# Immune experience at time of circulation and time since virus circulation are predictors of HAI titre

**DOI:** 10.1101/2023.07.25.550584

**Authors:** Joseph C. Gibson, Simon P. J. de Jong, Colin A. Russell

## Abstract

Nearly everyone is infected with seasonal influenza viruses multiple times over the course of their lives due to the antigenic evolution of the virus to escape immunity induced by prior infections and vaccinations. Because antibody escape is usually partial, the antibody response to new infections depends on prior responses which wane and are boosted throughout life. We used serum antibody binding measurements of 864 individuals against a range of historical A/H3N2 viruses collected in previous studies in combination with statistical models to investigate how metrics of age and strain heterogeneities affect haemagglutination inhibition titres. We refine prior modelling of antigenic seniority by characterising immune experience as years of life since a (sub)type’s emergence at time of strain circulation, rather than age at time of circulation. Using Bayesian statistical modeling, we show that this variable, combined with time since circulation of the variant and an individual’s age at circulation, yields the best model fit. Based on the best-fit model, we devised a novel parametric model, and use it to demonstrate and estimate how effects of antigenic seniority, individual age, strain age and strain effects act in concert to shape an individual’s HAI titre.

## 1 Introduction

Up to 30% of the world’s human population are infected with influenza each year [1], resulting in a substantial disease burden [2]. After infection with an influenza A virus (IAV), individuals develop some immune memory which reduces their probability of reinfection by similar antigenic variants. As a variant circulates and population immunity against it increases, viable hosts become progressively limited. In response to this immune pressure, the virus evolves to evade immunity, rendering enough of the population susceptible to ensure continued circulation. For influenza A/H3N2 viruses, this antigenic evolution results in a typical circulation period of 3-5 years for each new antigenic variant before it is replaced by the next one [3].

The head domain of the haemagglutinin (HA) surface protein forms the primary target for antibodies targeting influenza virus. Levels of HA head specific antibodies are typically characterised at the systemic level of the serum, by the ability of a blood sample to prevent viruses from binding to erythrocytes, known as the serum haemagglutination inhibition (HAI) titre. Serum antibody titres likely correspond to a high degree with those in the mucus, where virions may be intercepted at their points of entry and, indeed, HAI titres have been shown to offer a correlate of protection from infection [4, 5, 6].

Analyses of HAI titres to historical viruses have shown that an individual’s infection and subsequent periodical re-infection results in an antibody repertoire which is a complex mosaic of those previous infections. In particular, it has been shown that the first infections with a particular subtype take on a dominant position in one’s antibody repertoire, a notion referred to as original antigenic sin (OAS) [7]. The hierarchical boosting of cross-reactive memory responses upon subsequent reinfections with antigenically distinct viruses, which maintains the dominance of early-life infections, is referred to antigenic seniority (AS), which has been modelled as increasing suppression of HAI titre in a hierarchical manner with increasing age at time of exposure [8].

While past modelling endeavors, including those described above, have yielded crucial insights into the antibody dynamics of influenza virus throughout life, they rely on a crude dependence of titre on an individual’s age, which is potentially problematic as the emergence of antigenically novel influenza viruses can occur during one’s lifetime. Additionally, past models largely rely on spline fits to the data, impeding the estimation of biologically relevant parameters. Though cross sectional sera cannot provide a detailed individual infection history, data from China has been employed with success to elucidate population age patterns in titre [8]. Here, we make use of more recent data from this cohort [9] as well as previous cross sectional sera studies from Vietnam and Australia [10] totalling 864 individuals, where titres were measured against a range of historical A/H3N2 viruses. We employ statistical modelling to make refinements to the modelling of AS, and show that our modifications improve model fit. Additionally, we develop a novel parametric model for HAI titre data to estimate parameters of interest pertaining to AS, waning and strain effects.

## 2 Results

### Age at time of circulation imperfectly captures patterns of HAI titre

It has been previously demonstrated [8] that an individual’s microneutralisation titre against a particular strain depends on their age at time of circulation (AgeCirc), their age at time of sample collection (Age) and the strain, where titres were highest towards strains which circulated during the individual’s childhood, some weak age effects were present in the elderly and strain-to-strain variation was random. Whilst AgeCirc is a parameter which can encompass age dependent immune responses (for example, senescence) and exposure rates (for example, school attendance or vaccination rates), it cannot adequately serve as a proxy for *immune* experience as the H3N2 subtype only entered human circulation in 1968. For example, an individual born in 1940 will have been completely naive to H3N2, but not particularly young, during their first likely H3N2 exposure around 1968. Hence, we examined as a relevant variable how long an individual’s life has overlapped with the period of H3N2 circulation, which we dub the “Immune age at time of circulation” (ImmAgeCirc). Mathematically,

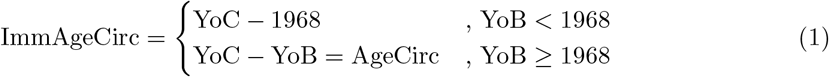

where YoC is the first year of virus isolation, which we assume to be the year of first circulation, and YoB is the year of birth.

To investigate if and how incorporation of this variable, as composed to simple age at strain circulation, might differentially explain patterns of HAI titre, we plotted titres for the China 2014 cohort against different values of age at time of circulation and *immune* age at time of circulation (Figure 1). For the strain that circulated in 1968, titre variation is largely absent (top left panel), despite values of AgeCirc ranging from 0 to 40 years which, according to previous work [8], should imply a titre drop of around 2 log units. Only the subset of the cohort with ImmAgeCirc *>* 0 sees titre variation by age, and this subset becomes progressively larger as year of strain circulation increases. Hence, it is only once later strains are considered so that ImmAgeCirc is able to vary do we see titre variation becoming progressively more pronounced. This is most evident for the 1982 strain (bottom right), where it could be argued that the variation in log titre for AgeCirc values 0-10 years is actually due to the changes in ImmAgeCirc. Hence, it is likely that patterns of titre variation are best captured by an individual’s *immune* age at time of strain circulation, rather than their simple age at time of circulation.

**Figure 1:**
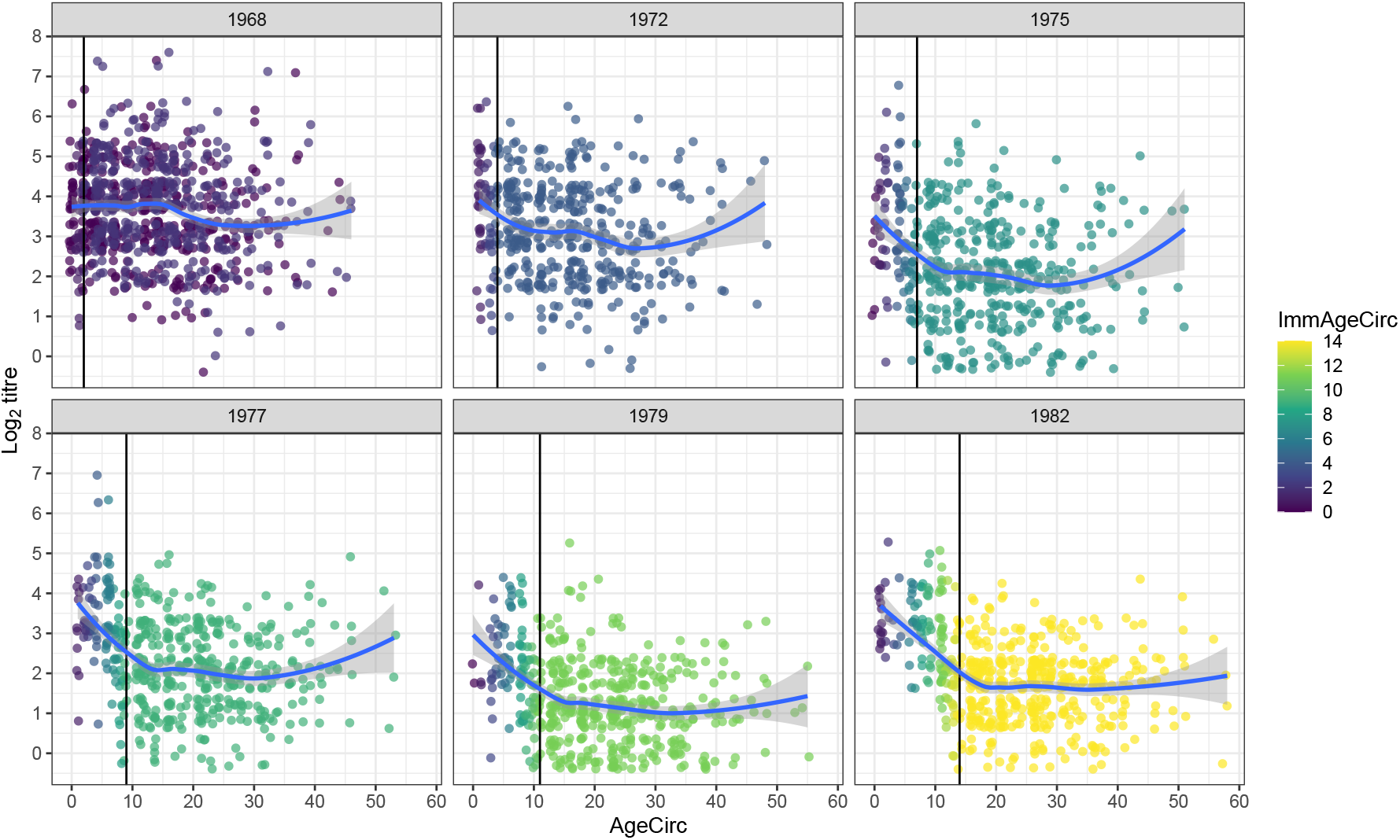
Variation in log titres for the China 2014 cohort with AgeCirc (x-axis) and ImmAgeCirc (colour) for early H3N2 strains. Points to the right of the black line indicate individuals born before 1968. Random noise has been added to data points and blue lines indicate a LOESS curve fit.

### Immune age at time of circulation and time since strain circulation best predict HAI titre

We tested the predictive value of ImmAgeCirc relative to AgeCirc more rigorously by fitting five generalised additive models (denoted A-E) to the four cross-sectional data sets. These models are formed by modelling log titre as the sum of strain effects plus combinations of splines of the parameters Age, AgeCirc and ImmAgeCirc:

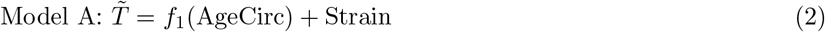

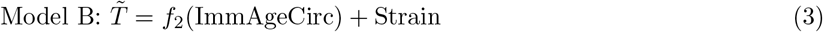

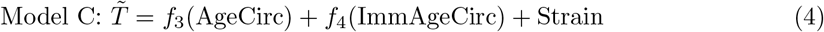

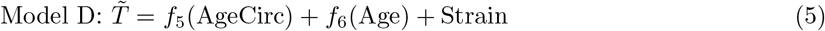

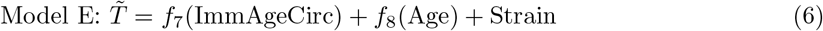

We note that Model D is the same as the one used in the original work on AS [8]. We fit these models to the data in a Bayesian framework, with a Markov Chain Monte Carlo (MCMC) algorithm used to explore the posterior parameter joint distribution.

To select the model that best explains the data from the fitted outputs, we computed differences between models in expected log pointwise predictive density (ELPD), a measure which rewards predictive capability whilst also penalising model complexity [11]. ELPD values are given relative to the best performing model and a more negative ELPD difference value indicates less evidence in favour of the model. Model C, which includes terms for AgeCirc, ImmAgeCirc and strain effects, was found to be the significantly best performer for the Australia 1997 and China 2014 data sets (Figure 2, top left and bottom right panels respectively). However, no outright superior model emerged for the Australia 1998 and Vietnam 2007 datasets, though in both cases the the best-performing model was Model E. Given the small differences in ELPD for the Vietnam data, it is likely that the confidence intervals predicted here are underestimates [12] and so no real conclusions can be drawn from this data set. The less clear-cut model selection for the Vietnam and Australia data sets is likely due to the minimal variation in birth year before 1968 and the limited spread outside the peaks of the bimodal age distribution of participants for the two countries, respectively (Supplementary Figure 2). Importantly, disentangling the relative effects of AgeCirc vs ImmAgeCirc on model fit requires a spread of birth years prior to 1968 to break the AgeCirc-ImmAgeCirc collinearity and a range of birth years post 1968 to break the ImmAgeCirc-Strain collinearity, respectively. Hence, we put more weight on the fact that Model C is the best predictor for China 2014, the largest and most uniformly distributed data set. It is also the most likely model for Australia 1997 and there is no strong evidence to reject it from the other two cohorts.

**Figure 2:**
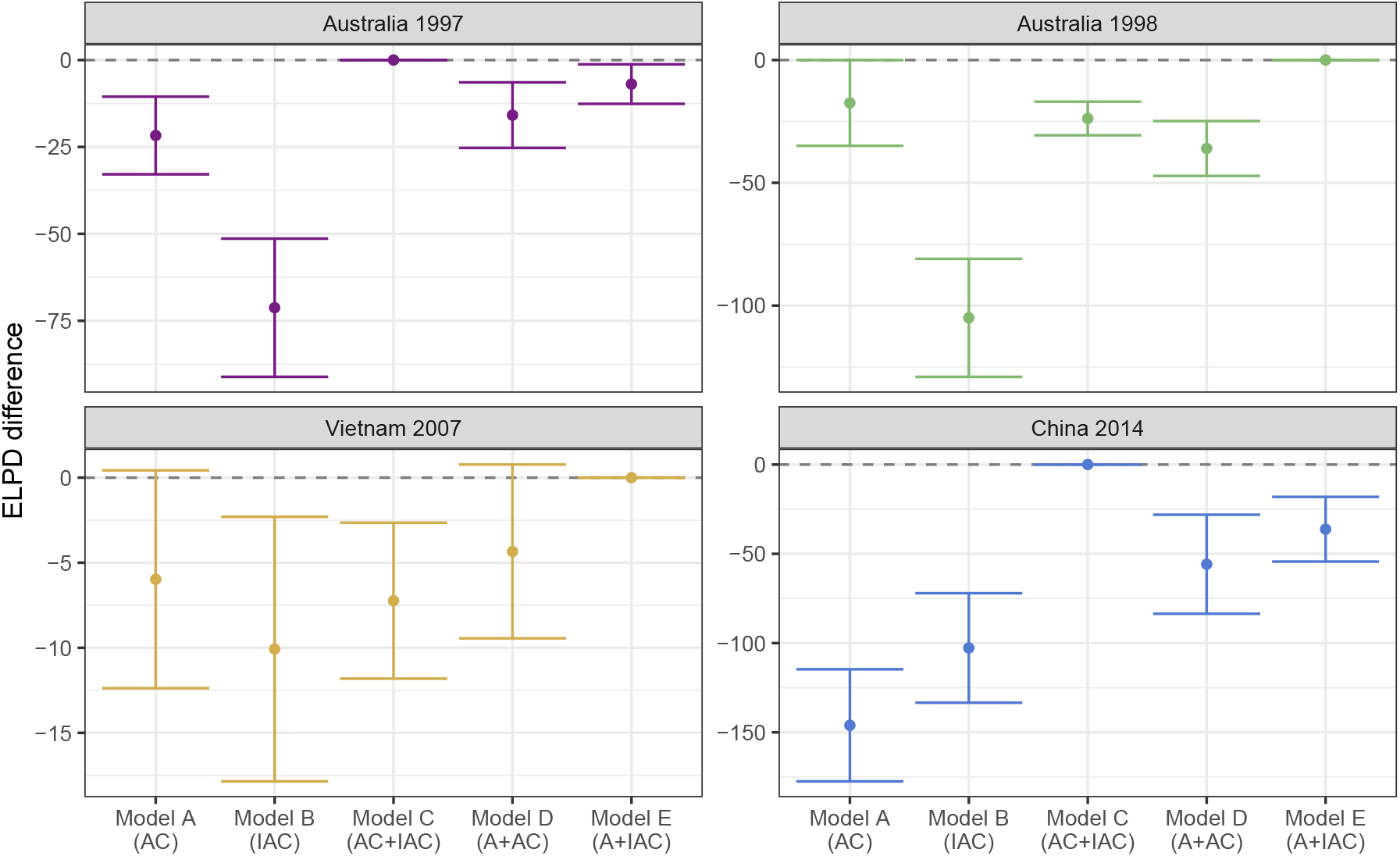
ELPD differences relative to the best performing model. 95% CI’s are shown, calculated as 1.96 times the estimated standard errors.

In the best-performing model, immunological inexperience at the time of strain circulation contributes significantly to titres being elevated (Fig 3a). This trend was quite consistent between data sets, with a decline of approximately 2-3 log units over a 30-40 year span of ImmAgeCirc. The AgeCirc spline (Figure 3b) was less well-matched between data sets, though for Australia 1997/8 and China the U-shaped curve indicates enhanced titres to strains encountered in childhood and older age, or perhaps suppression in middle age. Interestingly, thanks to the diversity in sample collection year between the data sets, we observed some suggestion that strain-specific effects are not entirely random. Rather, a trend appeared to exist with time since circulation (TSC) (Figure 3c), which can be seen by aligning the x-axis by sample collection year (Supplementary Figure 1). Strain intercepts generally followed a smooth pattern with year of circulation, whereby titres reach a peak around 10 years since sample collection and then decay over a period of a few decades. This would indicate that strain effects are not constant, but subject to change with time. In the next section we incorporate this time dependence into our modelling framework so as to investigate the likely underlying mechanisms.

**Figure 3:**
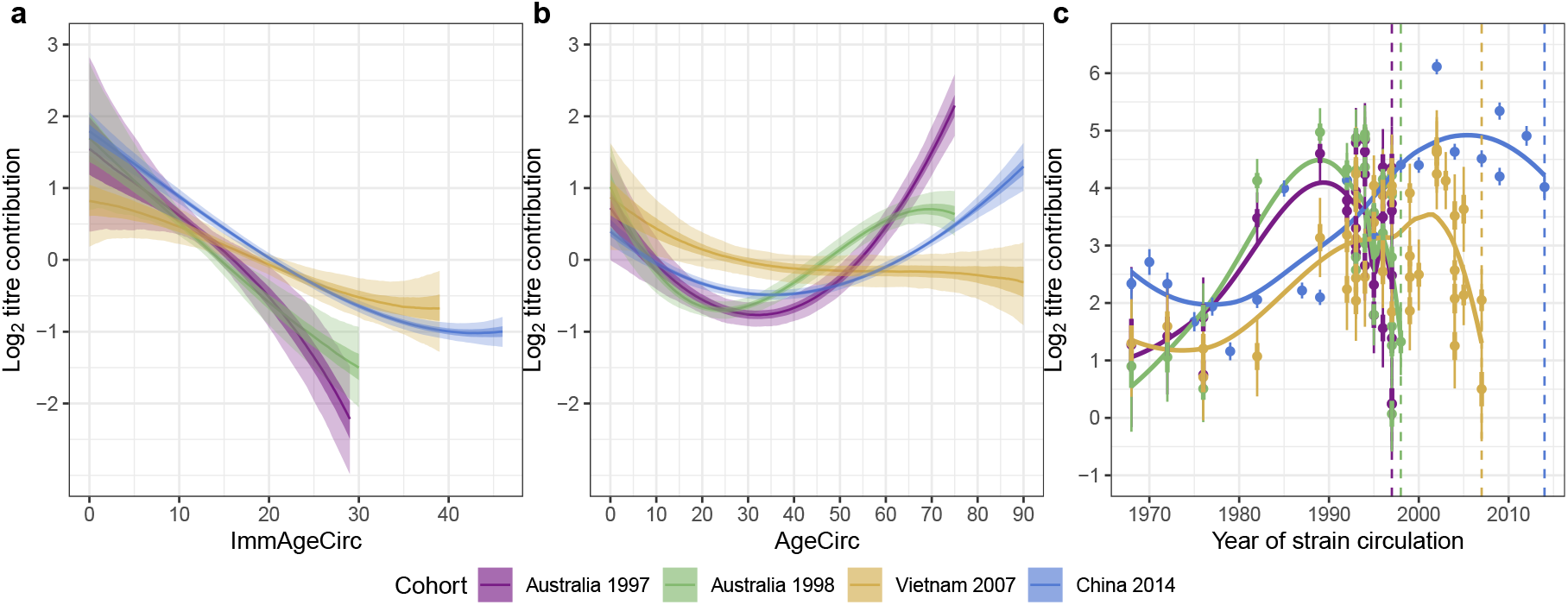
Splines and intercepts for the ImmAgeCirc (a), AgeCirc (b) and Strain (c) terms of Model C. Dark shaded areas in (a) and (b) and thick lines in (c) indicate 50% CI’s and light shading and thin lines indicate 95% CI’s. Vertical dashed lines in (c) indicate year of serum sample collection for each cohort.

### Parametric modelling of HAI titre

Our model fit exhibits some qualitative features that invite some biological and epidemiological interpretation (Figure 3). However, our model, following previous work on modelling of HAI titres, fits splines which are free to take any form, precluding the estimation of parameters of interest. To remedy this, we constructed a novel parametric model which constrains the fitting functions in order to gain estimates of parameters of interest, and includes the terms in the best-performing model in the model comparison analyses, and an additional term for time since circulation (TSC):

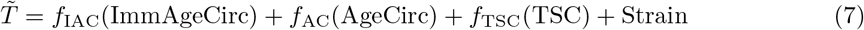

Briefly, we first assumed that ImmAgeCirc effects are described by a drop in titre of Δ*T*_AS_ between strains encountered when immunologically naive as compared to experienced, with this transition to experience occurring over a timescale of *τ*_AS_. Second, in absence of an obvious interpretation of the form of the AgeCirc curves, we observed that a parabolic function is sufficient to capture the shape of these profiles. Next, for TSC, we assumed two competing effects whereby titres build up over some timescale, *τ*_build_, but are also decreasing due to long-term antibody waning occurring at a rate *τ*_wane_. Lastly, we included random strain effects which are normally distributed with standard deviation, *σ*. This model is described in depth in the Methods by Eqs. (7)-(12), with parameter descriptions in Table 3. Supplementary Figure 3 schematically depicts this model, and illustrates how it parametrically reproduces the spline fits.

Given this novel parametric model, we used a Bayesian framework to estimate the parameters of interest, fitting again to the same four datasets. Titre drops due to AS were consistent across Australia 1997, Australia 1998, Vietnam 2007 and China 2014 at 3.26 (95% CI [2.07, 4.53]), 3.38 (95% CI [2.39, 4.3]), 1.96 (95% CI [1.43, 2.46]) and 2.53 (95% CI [2.27, 2.8]) log units respectively over a timescale of 22.3 (95% CI [17.1, 29.7]), 14 (95% CI [12.1, 16.7]), 14.4 (95% CI [11.5, 18.4]) and 19.2 (95% CI [17.5, 20.9]) years (Figure 4a and Table 1). For TSC, we observed a peak in titres to strains circulating around 5-10 years into the past with the long term dynamics of titres reflecting waning with half lives of 5.66 (95% CI [2.74, 12.2]), 4.06 (95% CI [2.34, 8.24]), 8.72 (95% CI [5.17, 13.6]) and 10.4 (95% CI [7.21, 14.9]) years. As we found in the previous spline fits, AgeCirc effects are quite different for the Vietnam cohort, though the large confidence interval indicates there is not enough data here to constrain the form well. Strain effects appeared to be random with a standard deviation ranging between 0.77 and 1.04 (Figure 4d). Large variation in strain effects for strains isolated in the same year was particularly noticeable in the Australia and Vietnam data. Strikingly, in all cohorts strain effects were higher than average in 1968 and then declined over the next few years, presumably a signature of high circulation after the initial introduction of the new subtype.

**Table 1:** Posterior distribution medians and 95% CIs (in square brackets) for the parameters characterising our second model.

| Parameter | Cohort |  |  |  |
| --- | --- | --- | --- | --- |
|  | Australia 1997 | Australia 1998 | Vietnam 2007 | China 2014 |
| a | 5.14 [3.77, 6.47] | 2.43 [1.45, 3.7] | 0.682 [0.067, 1.97] | 4.74 [4.03, 5.42] |
| b | 0.42 [0.341, 0.48] | 0.257 [0.044, 0.392] | 0.905 [0.313, 1.99] | 0.405 [0.378, 0.43] |
| $\Delta T_{AS}$ | 3.26 [2.07, 4.53] | 3.38 [2.39, 4.3] | 1.96 [1.43, 2.46] | 2.53 [2.27, 2.8] |
| $\tau_{AS}$ | 22.3 [17.1, 29.7] | 14 [12.1, 16.7] | 14.4 [11.5, 18.4] | 19.2 [17.5, 20.9] |
| $T_{peak}$ | 3.7 [1.45, 8.68] | 5.76 [2.98, 10.7] | 3.31 [1.82, 5.46] | 1.9 [0.319, 3.86] |
| $T_0$ | 1.81 [0.921, 2.66] | 1.37 [0.26, 2.42] | 1.42 [0.334, 2.63] | 3.55 [2.1, 4.71] |
| $\tau_{build}$ | 4.86 [0.994, 12.3] | 5 [1.87, 10.6] | 4.12 [1.58, 11.6] | 4.83 [0.439, 13.9] |
| $\tau_{wane}$ | 5.66 [2.74, 12.2] | 4.06 [2.34, 8.24] | 8.72 [5.17, 13.6] | 10.4 [7.21, 14.9] |

**Figure 4:**
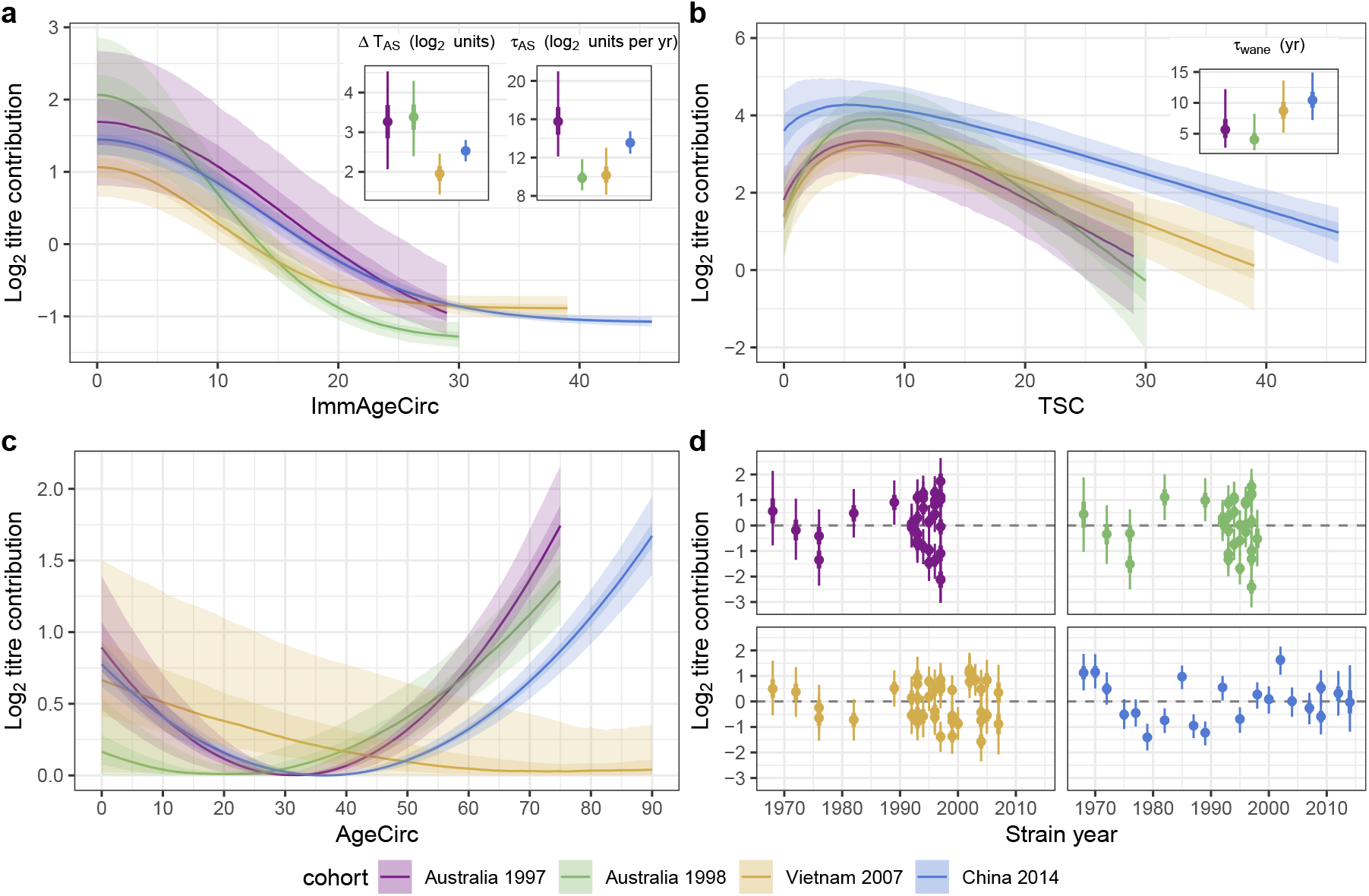
Model fit for the biologically based model. In (a), the expected ImmAgeCirc function with 95% CI and posterior distributions of key parameters *τ*_AS_ and Δ*T*_AS_ inset in the top right. In (b), the TSC function with 95% CI and posterior distributions of *τ*_wane_ inset. In (c) and (d) are the AgeCirc and random strain intercepts terms respectively. Points and lines indicate median values, dark shading and thick lines indicate 50% CI and light shading and thick lines indicates 95% CI which are derived from 1000 draws of the posterior distributions.

## 3 Discussion

In this study, we found evidence to support the inclusion of two previously overlooked variables which add power in predicting age dependent HAI antibody titres towards previously circulating H3N2 strains. These are the subtype specific years of experience at time of strain circulation (ImmAgeCirc) and the time since strain circulation (TSC). We found that these two variables, in tandem with the age at time of strain circulation (AgeCirc) and random strain intercepts gave the best performing model framework for predicting titres when judged by ELPD.

Our finding that ImmAgeCirc yields better model performance than AgeCirc is consistent with the prevailing mechanistic hypothesis of AS [13], whereby the first encounter with a completely unrecognised IAV results in antibody production towards a large fraction of HA epitopes. Then, in a subsequent infection with a new variant, activation of naive B-cells recognising the new epitopes is suppressed through antigen trapping by memory B cells, thus resulting in lower HAI titres than for the previous infecting strain. The time of exposure to the strain that attains the dominant positions in one’s antibody repertoire within this AS framework will necessarily be later than the time of emergence of that strain’s subtype. Hence, if an individual was born prior to the emergence of that subtype, AgeCirc is a flawed metric of immune experience in the context of antigenic seniority. We deigned a novel parametric model to be able to describe and quantify these effects, among others, expanding on the previously used modelling framework that relies on fitting splines. In this model, we introduced two parameters to provide a more rigorous characterisation antigenic seniority. We used *τ*_AS_, to capture the timescale over which repeated infections accumulate to suppress subsequent responses and found estimates to range from 14 to 22 years (Table 1). We also introduced Δ*T*_AS_ to quantify the drop in HAI titres resulting from a naive vs experienced infection and found estimates between 1.96 and 3.38 log units. This age threshold of immune maturity occurring in late teens could have particular significance in the context of describing how and when immune targets shift with age.

While we determined that ImmAgeCirc is an important variable when modelling titres, our best-performing model also included AgeCirc. We see the role of AgeCirc to have an effect in slightly enhancing titres for children and older adults, independent of ImmAgeCirc. This could be because children have higher exposure rates due to contacts at school and older adults could have higher immune stimulation due to increased vaccination rates. Whilst this latter reasoning might be applicable to the Australia data set, only 0.6% of the China data set reported vaccination between 2010 and 2014 and hence we cannot draw conclusions regarding the origin of this effect.

We also found time since strain circulation to be a predictor of titre. HAI titre increases for strains which circulated over the past decade or so, presumably due to increased exposure among the population at large. This population build up competes with individual waning such that a maximum is reached before waning takes over in the absence of strain circulation and thus immune stimulation. In our parametric model, we characterised this waning rate by *τ*_wane_ giving estimates for the antibody half life ranging from 4.06 to 10.4 years. This is an interesting parameter to characterise and, to our knowledge, has not been studied elsewhere. Whilst short term titre dynamics post infection have been well covered in the literature, long term waning of HAI has not, despite being quantified for a number of other viruses [14]. Our findings on the long term half life here are in agreement with estimates we extracted recently from two different longitudinal cohorts from the Netherlands [15]. Together, our findings add important new insight and nuance into the determinants of antibody dynamics to seasonal influenza viruses.

## 4 Methods

### Serology data

Our analysis is based on HAI titre data from sera collected from four different cohorts. Two are longitudinal in design, one being a study conducted in China [9], where samples were collected in 2010 with a follow up visit in 2014. The second is from Vietnam [10], where subjects gave samples on an annual basis between 2007 and 2012. Previous analysis of the Vietnam data in the context of antibody landscapes [10] has indicated likely infection events and these sample-patient years are omitted in our analysis to reduce any obfuscating effects of acute antibody dynamics following infection. The final two cohorts come from a vaccination study in Australia [10], involving two separate groups, one providing samples in 1997 and the other in 1998. We use the samples provided prior to receipt of the trial vaccine in our analysis, again to remove any obscuring effects of an active immune response. In all aforementioned studies, assays were performed using a selection of H3N2 strains isolated in a range of years from the introduction of H3N2 in 1968 to the year of sample collection.

Our main analysis is performed on China 2014, Vietnam 2007, Australia 1997 and 1998, so as to have four independent, temporally separated cohorts. We omit samples from strains which had not yet circulated at the time of sample collection since they are misleading with regard to the age at time of circulation. We also omit HAI samples from individuals towards strains which circulated before they were born, since there are important strain circulation effects which will be masked by those who were not alive at the time to be affected by them. The characteristics of each data set are outlined in Table 2. The age structure of the Australia data is highly dichotomous for both studies (Supplementary Figure 2a), with a lack of subjects born between 1940 and 1970. The Vietnam data is more uniformly spread but has few subjects overall, especially older ones. All studies cover a similar range of strain years, with the Australia and Vietnam data placing emphasis on those circulating after 1990 (Supplementary Figure 2b). It can be seen that by far the largest and most uniformly distributed data set is provided by the China cohort.

**Table 2:**
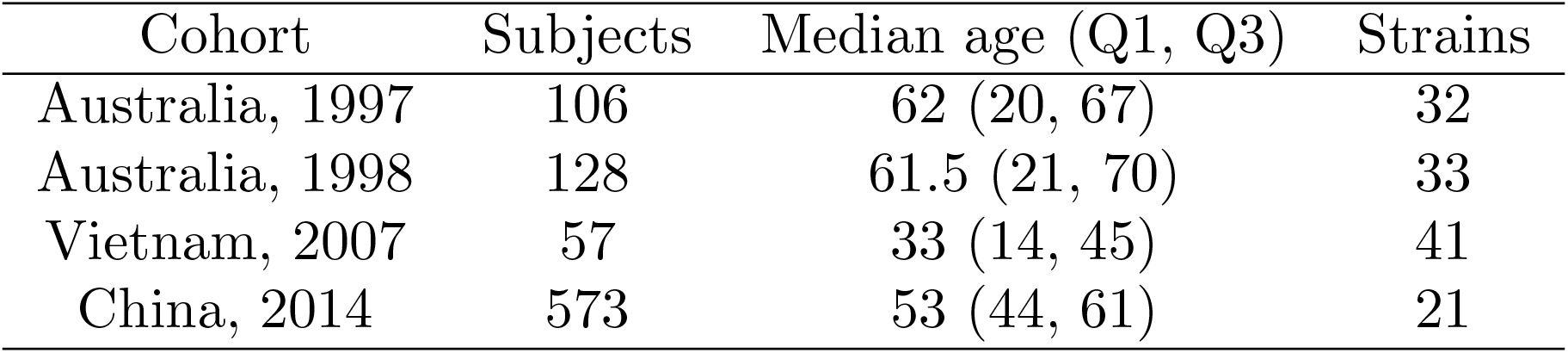
Characteristics of serum HAI titre data sets.

| Cohort | Subjects | Median age (Q1, Q3) | Strains |
| --- | --- | --- | --- |
| Australia, 1997 | 106 | 62 (20, 67) | 32 |
| Australia, 1998 | 128 | 61.5 (21, 70) | 33 |
| Vietnam, 2007 | 57 | 33 (14, 45) | 41 |
| China, 2014 | 573 | 53 (44, 61) | 21 |

### Titre modelling

In modelling titre, each of our candidate models predicts mean true titre, 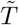, as a function of variables, **x**, (Strain plus some combination of AgeCirc, ImmAgeCirc and Age) and parameters ***θ***. To compare with observations, we must include a measurement model, which gives the likelihood of making a certain measurement, *T*, given the expected value, 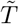, measurement error and individual variation. This measurement model reflects the fact that titre measurements are made in discrete dilution increments and are conducted between maximum and minimum concentrations of 8 and 1 rounds of dilution respectively.

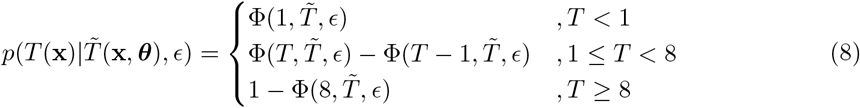

where Φ(*x, μ, ϵ*) is the cumulative distribution function of the normal distribution with mean, *μ*, and standard deviation, *ϵ*. Measurement error and individual variation are assumed normally distributed and can both be encapsulated by a single standard deviation, *ϵ*. The total likelihood is thus the sum of Equation (8) over all data points.

### Spline model

In the first analysis, log titre is modelled to have a variety of possible dependences through the five candidate generalised additive models described in the main text. In those equations, the unknown functions *f*_1*−*8_, are formed by a superposition of third degree B-splines, implemented in R v4.0.3 using the splines2 v0.4.5 package.

### Parametric model

In the second analysis, after finding model C to be the most likely choice, we test explicit forms for the functions in Equation (4) so that we may extract some biological insight. The explicit form for the equations describing each of the constituent effects are as follows:

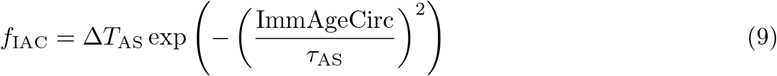

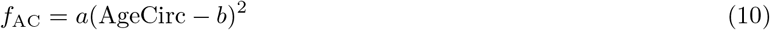

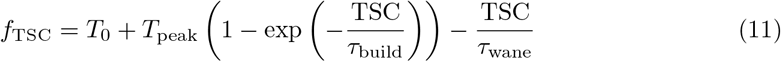

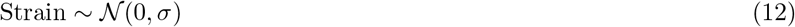

The descriptions of the parameters of Equtions (7)-(12) are given in Table 3, as well as a schematic diagram in Supplementary Figure 3 to aid in their interpretation.

**Table 3:**
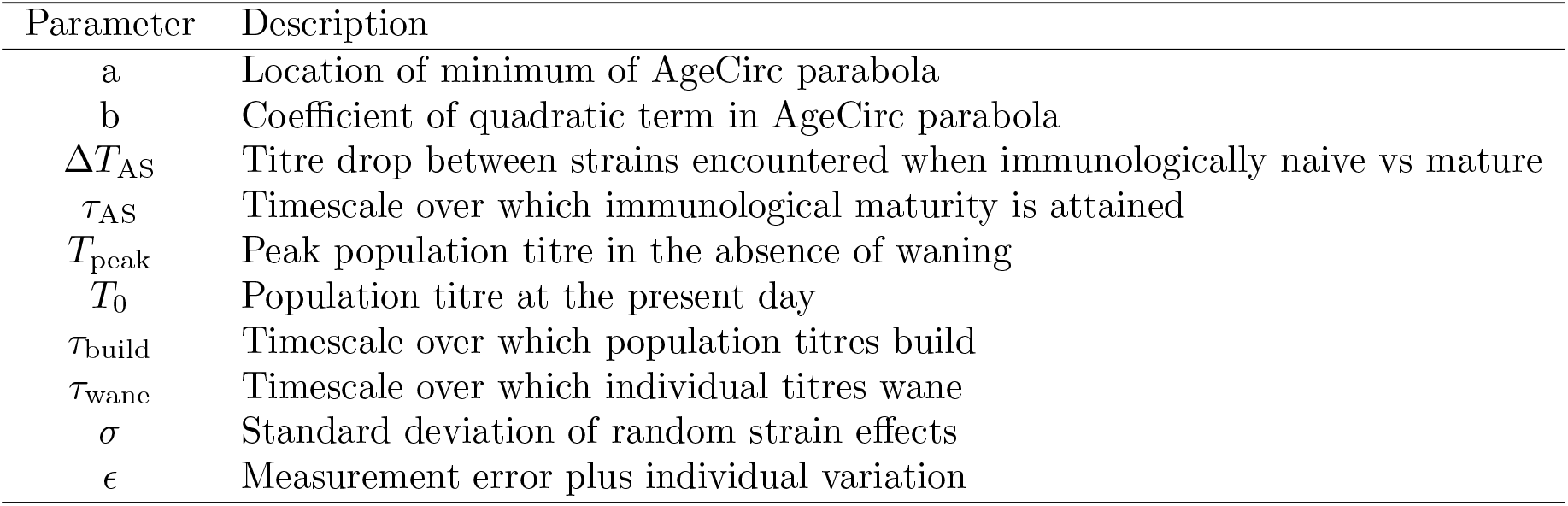
Description of parameters used in our biologically based model.

| Parameter | Description |
| --- | --- |
| $a$ | Location of minimum of AgeCirc parabola |
| $b$ | Coefficient of quadratic term in AgeCirc parabola |
| $\Delta T_{AS}$ | Titre drop between strains encountered when immunologically naive vs mature |
| $\tau_{AS}$ | Timescale over which immunological maturity is attained |
| $T_{peak}$ | Peak population titre in the absence of waning |
| $T_0$ | Population titre at the present day |
| $\tau_{build}$ | Timescale over which population titres build |
| $\tau_{wane}$ | Timescale over which individual titres wane |
| $\sigma$ | Standard deviation of random strain effects |
| $\epsilon$ | Measurement error plus individual variation |

A Markov Chain Monte Carlo (MCMC) algorithm was used to explore the joint distribution of model parameters. This was run on four independent chains, each consisting of 5000 iterations with the first 2500 discarded as burn in. Weakly informative priors were used:

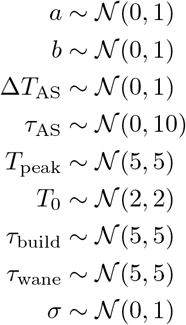

and convergence was assessed by inspection of the trace plots and Rhat from Stan v2.21.0. Analyses were conducted using R v4.0.3.

## 5 Acknowledgements

This work was supported by grants from the Wellcome Trust (200187/Z/15/Z) and the European Research Council (818353)

