## Supplementary Figures for "Immune experience at time of circulation and time since virus circulation are predictors of HAI titre"

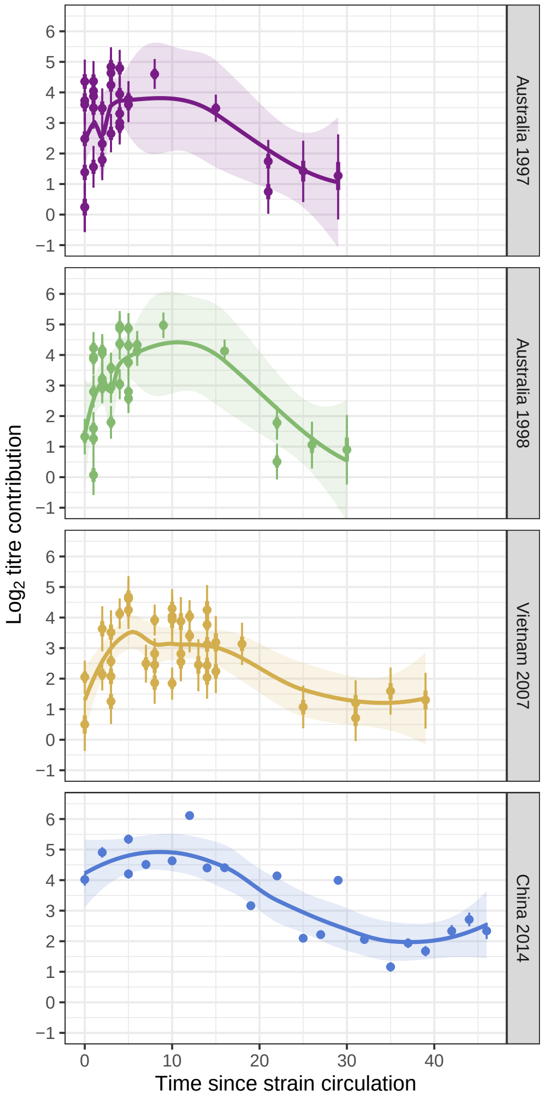


Supplementary Figure 1: Strain intercepts of Figure 3 aligned by sample collection and plotted against TSC. Lines indicate a LOESS curve fit.

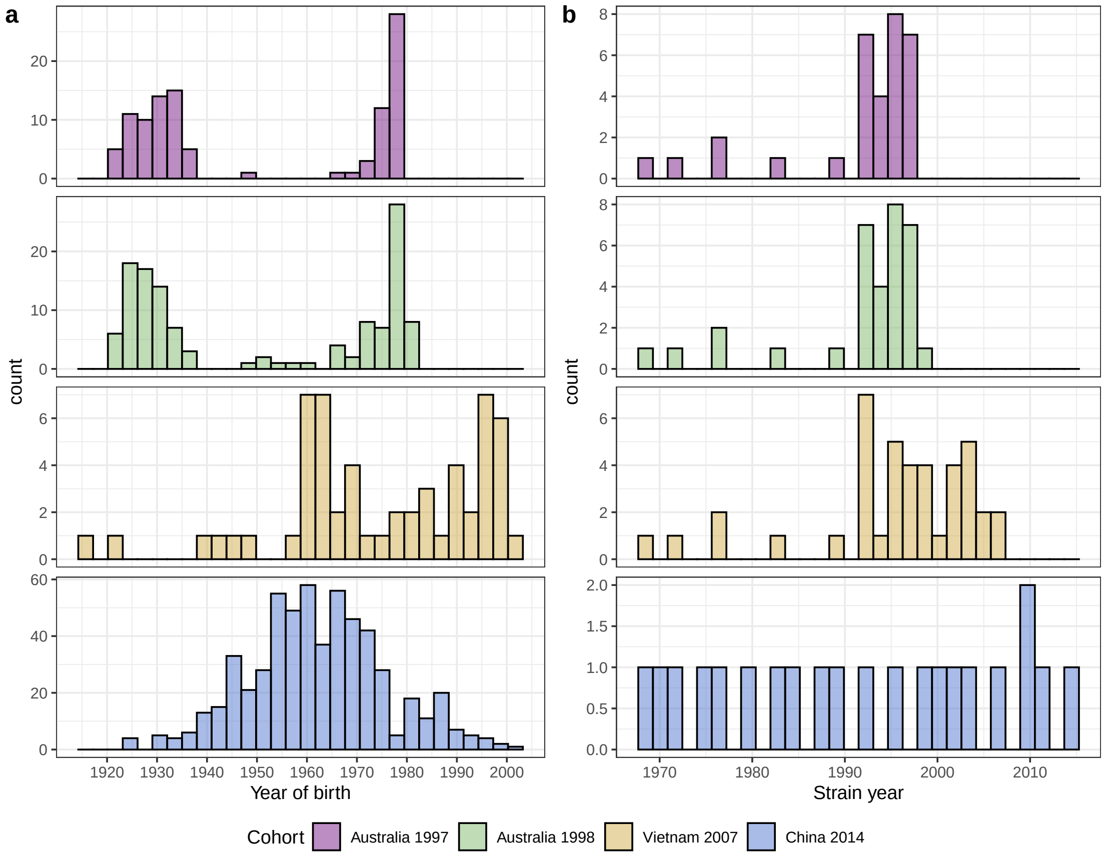


Supplementary Figure 2: Distributions of participant year of birth (a) and year of first isolation of the assay strains used (b) for the four HAI data sets.

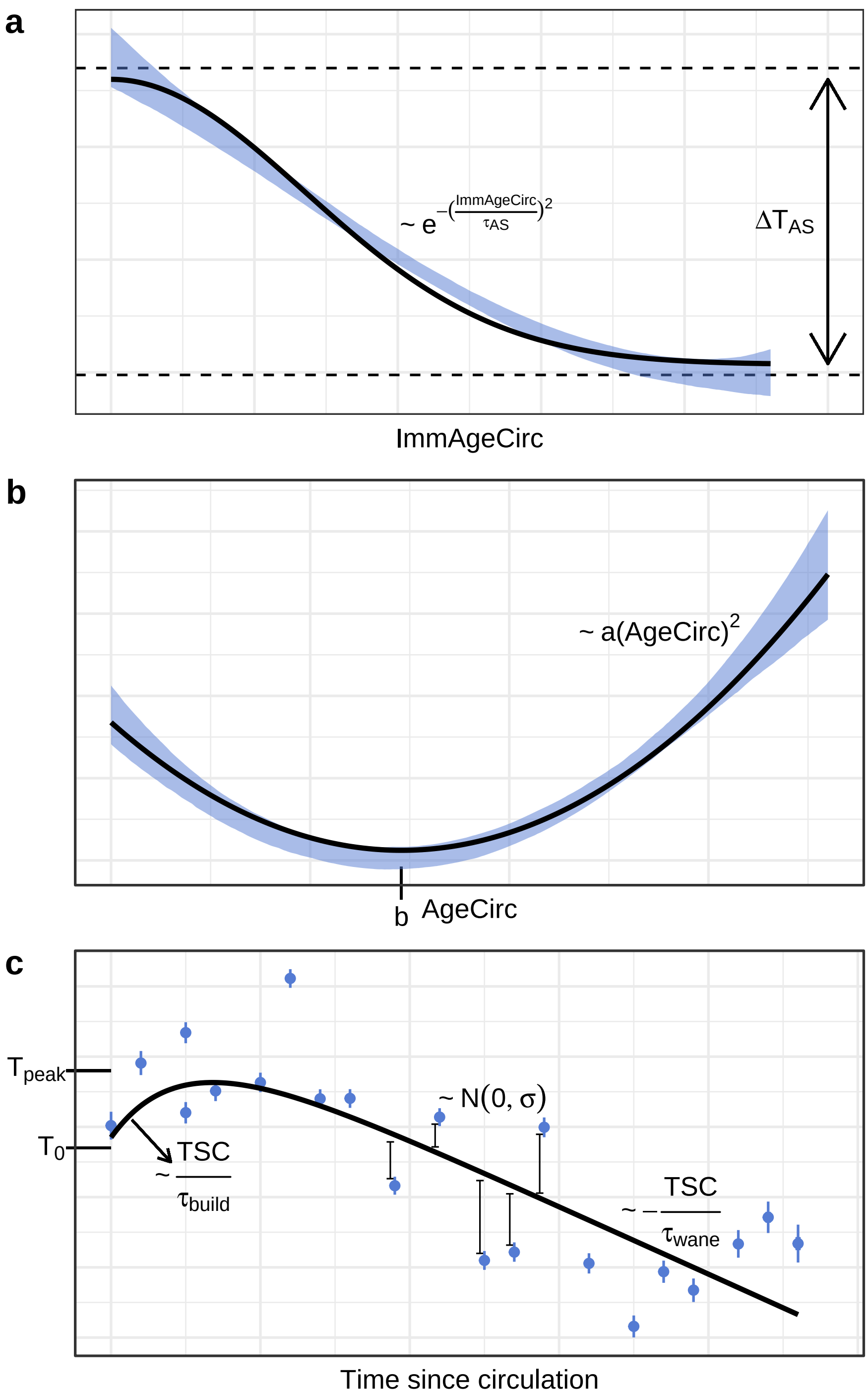


Supplementary Figure 3: Illustration of Equations (9)-(11) in panels (a)-(c) respectively. The blue areas in (a) and (b) correspond to the 95% CI of the spline fits for the China 2014 cohort in Figure 3 and the blue points in (c) to the strain intercepts. Black lines correspond to the trial functions in Equations (9)-(11) where the maximum likelihood parameters have been used.
